# Dynamics and Determinants of the Gut Mobilome in Early Life

**DOI:** 10.1101/2024.09.06.611692

**Authors:** Asier Fernández-Pato, Trishla Sinha, Sanzhima Garmaeva, Anastasia Gulyaeva, Nataliia Kuzub, Simon Roux, Jingyuan Fu, Alexander Kurilshikov, Alexandra Zhernakova

## Abstract

As mobile genetic elements (MGE) are critical yet understudied determinants of gut microbiome composition, we characterized the gut virome and plasmidome in 195 samples from 28 mother–infant dyads delivered by caesarean section. Infant mobilome increased in richness over the first 6 postnatal weeks, demonstrating high individual-specificity and temporal stability, establishing a personal persistent mobilome. Formula-fed infants exhibited greater mobilome richness than breastfed infants, with plasmid composition influenced by antibiotic exposure and birth weight. Plasmids constituted a significant reservoir of antibiotic resistance genes (ARG), with around 5% of infant gut plasmid taxonomic units carrying ARG. Notably, ARG profiles did not differ with antibiotic exposure at birth. We found that mother–infant sharing of viral and plasmid strains primarily occurred after 6 months of age. Overall, our integrative analysis offers novel insights into the dynamics, modulation, origin, and clinical implications of MGE in the developing gut microbiome.

## Introduction

In recent years, research into the early-life human gut microbiome has revealed a complex ecosystem with the potential to influence immune development and disease susceptibility at later stages of life.^1–3^ Among the factors influencing the composition and function of the gut microbiota, mobile genetic elements (MGEs) have emerged as key players in shaping microbial evolution.^4–6^ MGEs, including plasmids, transposons, and bacteriophages, possess the remarkable ability to transfer genetic material horizontally between bacterial species, thereby driving genetic diversity and adaptation within microbial communities.^7–12^ MGEs are of particular interest given their potential role in disseminating antibiotic resistance genes (ARGs) within the gut microbiota through horizontal gene transfer (HGT).^9,10,13–15^

Advances in high-throughput sequencing enable exploration of the total DNA present in environmental samples, including the human gut. Recently, metagenomic assembly approaches have been applied to reconstruct infant gut viral communities (virome) from shotgun sequencing data.^16–22^ These early-life studies have identified associations with clinical factors such as delivery mode, gestational age, and feeding patterns,^19,21^ as well as links to conditions like asthma.^22^ However, research on the infant plasmid community (plasmidome) remains limited to one recent large-scale study.^23^ Therefore, a critical gap exists in longitudinal studies exploring the co-development of the gut virome and plasmidome in early infancy. This is particularly relevant in the first weeks of life as infant gut microbial colonization occurs rapidly after birth and any disturbance in this process may influence later development of diseases.^1,24–26^ Understanding the dynamics of MGEs during this critical window, as well as the forces shaping their diversity and composition, is essential for elucidating their impact on the establishment of microbial communities and the emergence of antimicrobial resistance.

The sources of ARGs in infants also remain unclear. It has been hypothesized that different factors, including mode of delivery, gestational age, infant feeding mode (breast milk or formula milk), and the mother’s previous antibiotic exposure, might influence the infant’s antibiotic resistance profile in early life.^27–29^ During caesarean sections (CS), pregnant women usually receive antibiotics pre-incision as a prophylactic method against infection. Newborns delivered by CS consequently exhibit prenatal antibiotic exposure via the umbilical cord due to the rapid placental transfer of these medications, and we previously showed this to have subtle effects on bacterial strain profiles.^30^ Similarly, earlier evidence suggests a potential maternal origin of the MGEs colonizing the early infant gut, with increased MGE presence in the infant potentially facilitating transmission of ARGs across microbial species.^29,31,32^ However, few studies have explored the effect of the prophylactic antibiotics administered prior to birth on the infant resistome and MGE-mediated maternal-offspring ARG transmission.

In this study, we explored longitudinal metagenomic sequencing data from infants born via CS as part of the CS Baby Biome randomized clinical trial (RCT). First, we comprehensively characterized the dynamics of the infant gut viral and plasmid communities during the first 6 weeks of life. We then explored what factors influenced the diversity and composition of infant MGEs, including the impact of prophylactic antibiotic timing at birth and infant feeding mode, as well as other pregnancy, delivery, and environmental factors. Finally, to understand the clinical implications, we investigated the role of MGEs as reservoirs for ARGs during early life and potential mother-to-infant MGE transmission.

## Results

### Characterization of the early infant gut viral catalogue

Using 195 faecal samples collected from infants and mothers from the CS Baby Biome cohort,^30^ we first aimed to characterize the double-stranded DNA viruses present in the infant gut during the first 6 weeks of life. In total, we longitudinally followed 28 infants born by CS at six timepoints and characterized their mothers at one timepoint collected around birth. In addition to the biological specimens, we had phenotypic information collected from all participants (see STAR Methods) (Table S1). Using an assembly-based virus identification framework, we identified 2,263 viral operational taxonomic units (vOTUs) that were present in at least one of our samples, including 630 specific to our dataset, i.e. vOTUs not clustering with any viral genome from other databases (Figure S1A). Nearly half of the vOTUs (48.9%) were singletons, meaning they consisted of a single viral genome (n=1,108) (Figure S1B). Regarding the genome quality of the identified viral species: 216 vOTU representatives (9.5%) were identified as complete viral genomes, 1,105 (48.9%) were classified as high-quality genomes, and 861 (38%) were classified as medium-quality genomes (Figure S1C). The taxonomic assignment of the vOTUs showed that the dsDNA human gut virome was largely dominated by *Caudoviricetes* phages, which were 96.5% of the viral species identified (Figure S1D), and most vOTUs were predicted to be temperate viruses (82.5% of the assigned vOTUs, Figure S1E). We then clustered vOTU-representative sequences into higher-ranking viral clades using pairwise average amino acid identity (AAI) and gene-sharing profiles.^33^ This yielded 699 genus-level and 245 family-level vOTUs, of which 147 and 46, respectively, were exclusive to the CS Baby Biome dataset, thereby enhancing the known early-life gut viral diversity. These findings underscore the importance of assembly-based viral discovery and catalogue generation, as the infant gut virome remains underrepresented in existing public databases.

### The infant gut virome is stable, persistent, and individual-specific over the first 6 weeks of life

Alpha diversity analysis of infant vOTUs did not reveal significant differences in the first 6 weeks of life (Figure 1A) (linear mixed-model, β=0.02, p=0.53). Similarly, we found that the proportion of predicted temperate phages did not differ between timepoints (Figure 1B) (linear mixed-model, β =-0.51, p=0.23). In contrast, we found that viral species richness increased over time (Figure 1C) (linear mixed-model, β = 1.37, p=9.08e-06), with the infant gut at 5 and 6 weeks of life showing a significantly higher number of species than the infant gut at week 1. Overall virome composition did not show a significant association with time (Figure 1D) (linear mixed-model, p=0.41). We further compared infant samples from the first 6 weeks of life with available maternal samples (n=23) in which we identified 1,174 vOTUs. This revealed a distinct gut virome profile between mothers and infants, with maternal samples showing higher diversity (Figure S2A, S2B) (linear mixed-model, β =1.70, p=2.34e-19) and a lower proportion and relative abundance of temperate phages (Figure S2C, S2D) (linear mixed-model, β =-21.27, p=2.15e-13 (proportion); β =-29.41, p=2.15e-8 (relative abundance)). Overall, the maternal gut showed a distinct virome composition in comparison to the early infant gut (Figure S2E) (linear mixed-model, p=6.2e-4), consistent with previous studies.^16,17,19^

**Figure 1.**
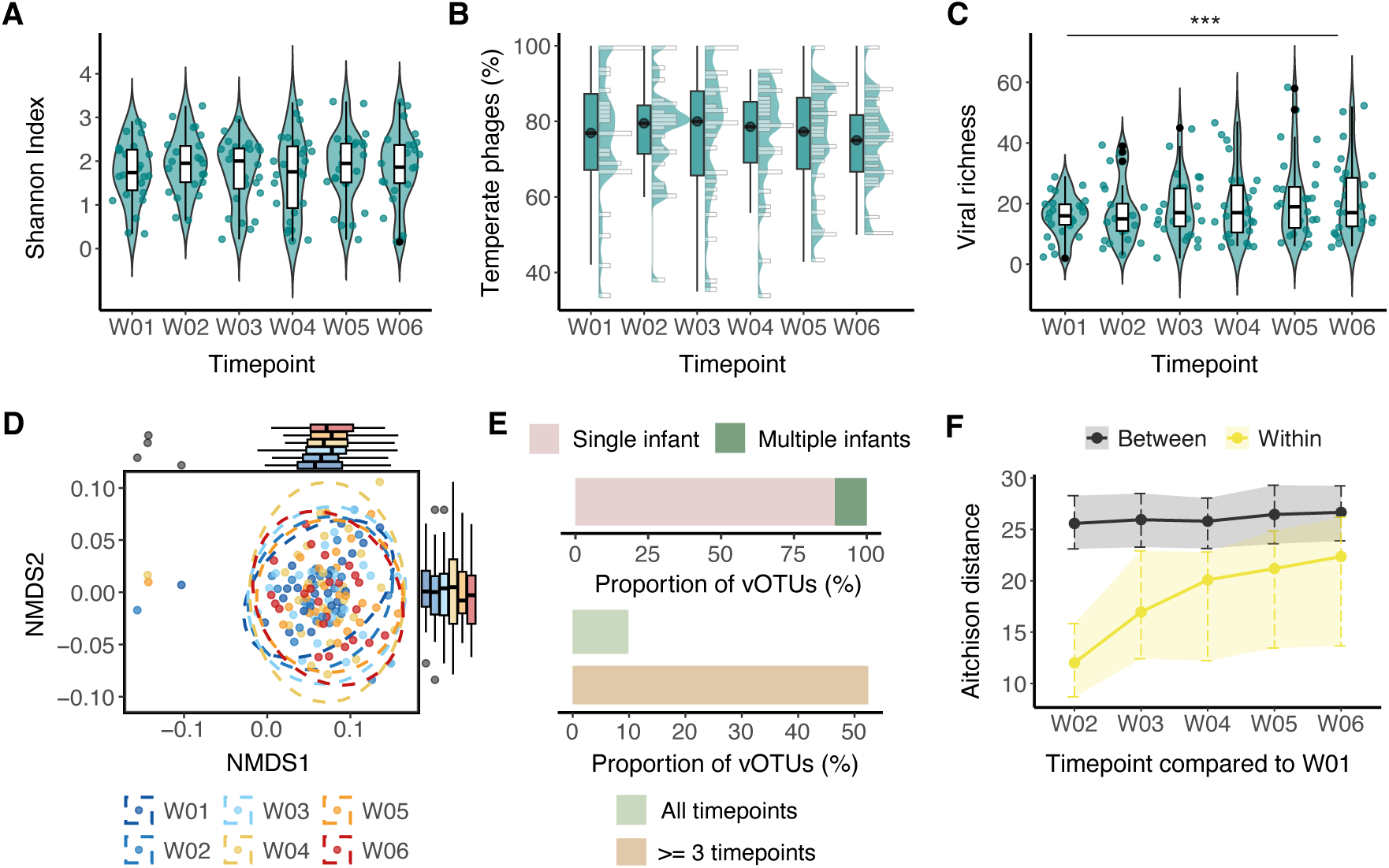
The infant gut virome is stable, persistent, and individual-specific over the first 6 weeks of life. (A) Alpha diversity (measured as Shannon Index), (B) proportion of temperate phages, and (C) vOTU richness in infant samples over the first 6 weeks of life. (D) Nonmetric multidimensional scaling (NMDS) plot based on Bray-Curtis dissimilarity estimated on vOTU abundances. Each dot represents a sample, coloured according to timepoint. Ellipses represent the 95% confidence regions for each timepoint, assuming a multivariate t-distribution of the data points. In all cases, box plots show the median values (middle line), interquartile range (box boundaries), and 1.5 times the interquartile range (whiskers). (E) Bar plots illustrating the proportion of all infant vOTUs found in a single infant (pink) versus multiple infants (green) and the proportion of vOTUs present in at least three timepoints (light brown) or all timepoints (light green) within the same infant over the first 6 weeks of life. (F) Dot plot showing the centroids of Aitchison distances between samples from each timepoint and week 1. Distances within the same infant are shown in yellow. Distances between different infants are shown in black. The interquartile range (IQR) for each comparison is also displayed.

We next explored the individual-specificity and temporal stability of the early infant gut virome (weeks 1–6). We observed that a low proportion of vOTUs were detected in multiple infants, with 89% of the vOTUs (n=830) found only in a single individual.

However, 52.4% (n=489) of the vOTUs were detected in at least three timepoints of the same infant (persistent vOTUs), with 9.9% (n=92) persistently present through all 6 weeks of early life (Figure 1E). Given that previous studies have suggested a role for diversity-generating retroelements (DGRs) in phage persistence in the gut through promotion of bacterial host switching,^20,34^ we investigated the potential impact of DGRs in our infant gut vOTUs. We found that DGRs were rare in early-life vOTUs (n=11, 1.2%) and significantly less frequent than in maternal vOTUs (n=194, 16.5%) (Chi-squared test, p=9.54e-32) (Figure S2F). Analysis of persistent vOTUs revealed no association between the presence of DGRs and vOTU persistence in infants (Fisher’s exact test, p=0.55).

We also compared the within- and between-individual Aitchison distances based on vOTU abundances over weeks 1–6. This analysis demonstrated consistently lower within-infant distances compared to distances between different infants, at all timepoints, including the samples taken 6 weeks apart (Figure 1F) ((Permutation-based p < 0.001). Altogether, these results demonstrate the infant gut virome’s diversity and compositional stability, as well as its pronounced individual-specificity and persistence in the first weeks of life.

### Feeding mode, rather than antibiotic timing at birth, shapes the early infant gut virome

To identify factors that influence the gut virome composition in the developing infant gut, we explored the associations between available metadata (see Methods) and virome features including alpha and beta diversity and viral abundances. Our analysis, which was adjusted for infant age, revealed significant associations between viral richness and infant feeding mode (linear mixed-model, β=14.12, FDR-adjusted p=4.25e-3) and the presence of pets in the family (linear mixed-model, β=12.85, FDR-adjusted p=1.87e-2). However, the presence of pets was no longer significant after also adjusting for feeding mode. The overall virome composition and the relative abundance of temperate phages did not show any significant associations with any metadata (Table S2).

Since the high individual-specificity of the vOTUs in the early gut virome represents a challenge when attempting to find associations between individual vOTUs and metadata, we performed our association analysis by grouping vOTU abundances according to their predicted bacterial hosts. In total, 94.2% of all vOTUs could be assigned a bacterial host at genus-level, with 53% of the prevalent vOTUs showing a positive correlation with their host’s abundance (Spearman correlation, FDR-adjusted p<0.05) (Table S3). This approach, adjusted for infant age, identified a positive association between the abundance of phages predicted to infect *Clostridium* and *Lacticaseibacillus* and infant formula-feeding (linear mixed-model, FDR-adjusted p<0.05). We also observed that the abundances of *Enterococcus_D* and *Limosilactobacillus* phages were positively associated with maternal pre-pregnancy BMI and the presence of household pets (Table S4).

Given the high correlation between viral abundances clustered by the bacterial host and the bacterial abundances, we investigated which associations persisted after adjusting for host abundance. Notably, none of the associations remained significant after FDR-correction, suggesting that changes in phage abundances are mainly driven by changes in the abundance of their bacterial host (Table S5). Importantly, none of our analyses revealed significant associations of infant gut virome features with the timing of antibiotic treatment at birth. All these findings highlight infant feeding as a significant factor shaping the early infant gut virome, largely driven by changes in the abundance of bacterial hosts, while the timing of antibiotic administration to the mother during CS birth appears to have a minimal or insignificant effect.

### Expanding the known plasmid diversity in the developing infant gut

Beyond gut phages, plasmids are also crucial in determining bacterial fitness through the dissemination of cargo genes. To better understand plasmids’ significance in the developing infant gut, we characterized the plasmid community by reconstructing and identifying both circular plasmids and plasmid fragments in our metagenomic samples. We detected 3,477 plasmid taxonomic units (PTUs), with 1,533 present in samples from the first 6 weeks of life (early-life PTUs). Among all PTUs, 36.6% (n=1,273) were circular (Figure 2A). The majority of PTUs were classified as non-mobilizable (53%, n=1,842), but smaller yet significant fractions were identified as mobilizable (40%, n=1,393) or conjugative (7%, n=242) (Figure 2B). Circular PTUs exhibited a greater length compared to plasmid fragments (Figure S3A) (one-sided Wilcoxon rank sum test, p<0.001). As the method we used to identify plasmid mobilization potential (see Methods) relies on the completeness of the plasmid genome, mobilizable (42.3%, n=539) and conjugative (9.6%, n=122) PTUs were more prevalent within the circular plasmid population. Nonetheless, we still found complete conjugative systems in 5.4% of the fragmented PTUs (Figure 2C). Conjugative PTUs were also more prevalent and displayed a significantly increased mean abundance compared to mobilizable (post-hoc Dunn’s test, FDR-adjusted p=4.68e-4 for prevalence; FDR-adjusted p=8.58e-10 for abundance) and non-mobilizable PTUs (post-hoc Dunn’s test, FDR-adjusted p=4.61e-5 for prevalence; FDR-adjusted p=2.39e-16 for abundance) (Figure 2D, Figure S3B).

**Figure 2.**
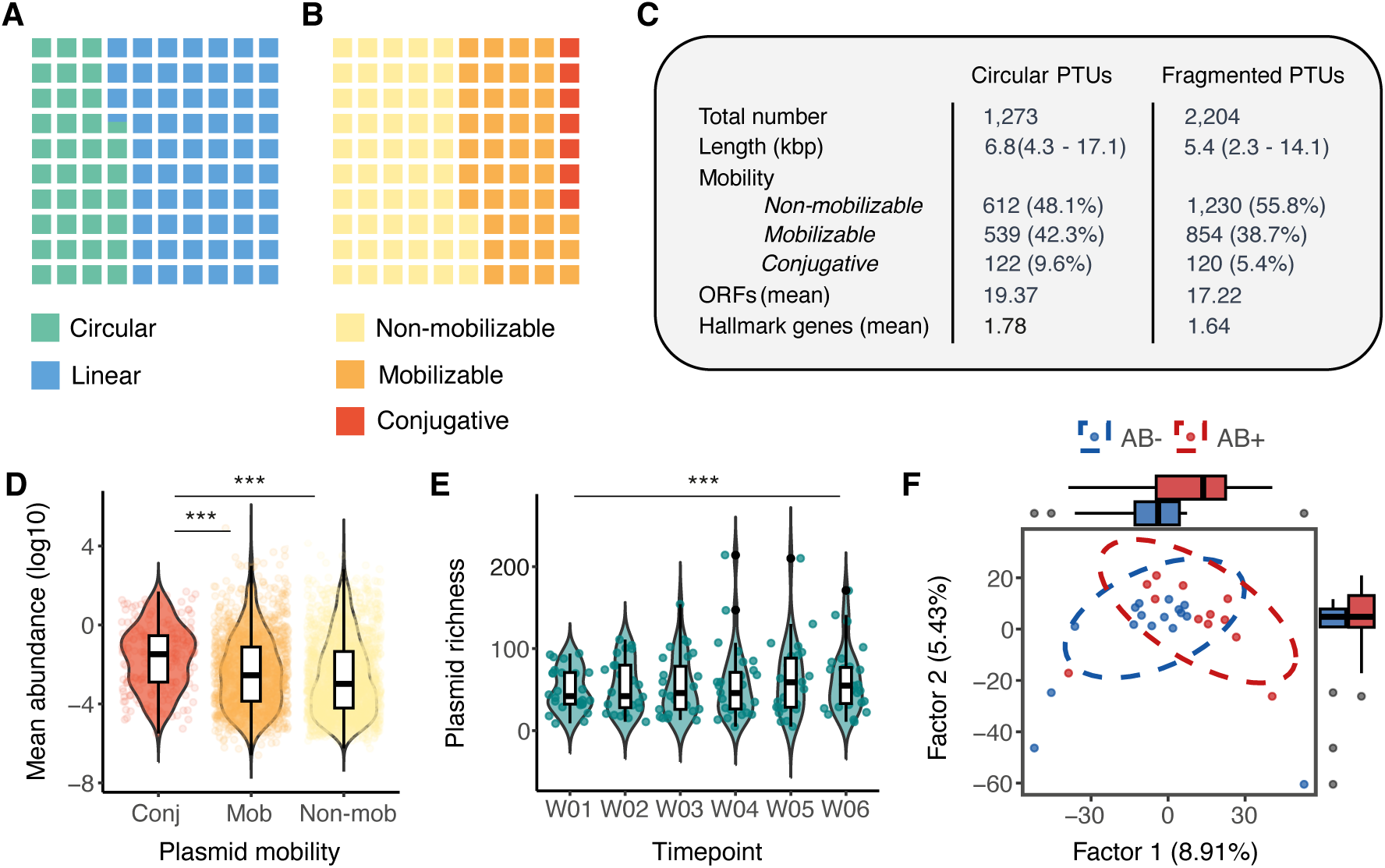
Characterization of plasmid community and its influencing factors in the developing infant gut. (A,B) Waffle charts of the proportion of gut PTUs classified according to (A) their structure, circular (green) or linear (blue), and (B) their mobilization potential, divided into non-mobilizable (yellow), mobilizable (orange), and conjugative (red) categories.(C) Summary statistics of the PTUs, including total number, genome length (in kilobases), distribution by mobility group, average number of predicted open reading frames, and average number of plasmid hallmark genes. (D) Boxplots and violin plots displaying the PTU mean abundance (log10 scale) according to the plasmid mobility. (E) Plasmid richness in infant samples over the first 6 weeks of life. Box plots show the median values (middle line), interquartile range (box boundaries), and 1.5 times the interquartile range (whiskers). (F) Scatter plot of the leading TCAM factors estimated based on PTU abundances in infant samples. Each dot represents the PTU composition of a single infant over the first 6 weeks of life. Colours represent the infants’ exposure to prophylactic antibiotics at birth. Blue indicates the non-exposed group (antibiotics administered after umbilical cord clamping). Red indicates the exposed group (antibiotics administered before skin incision). Ellipses represent 95% confidence regions of each group, assuming a multivariate t-distribution of the data points. Box plots for both TCAM factors are also shown.

Focusing on early-life PTUs, we observed that 36.5% (n=560) were found in multiple infants. Moreover, PTUs exhibited high temporal stability, with 15% of PTUs consistently detected across the first 6 weeks in the same infant (Figure S3C). Notably, we found that plasmid mobilization potential and circularity were associated with PTU persistence in the first 6 weeks (Chi-square test, p=1.79e-4 for mobilization potential; p=2.01e-8 for circularity), with conjugative and circular plasmids displaying increased persistence. Overall, our infant gut plasmid catalogue offers a comprehensive view of the early gut plasmid community, emphasizing conjugative potential as a key factor in determining plasmid persistence and abundance in the developing infant gut.

### Infant feeding, birth weight, and antibiotic timing at birth affect the early gut plasmid diversity and composition

Similar to viral community results, we found that overall plasmid richness and alpha diversity was significantly higher in mothers compared to infants (linear mixed-model, β=421.0, p=2.69e-72 for richness; β=2.51, p=1.30e-38 for alpha diversity) (Figure S3D, S3E). We also observed that plasmid richness increased slightly during the first 6 weeks of life (linear mixed-model, β=2.48, FDR-adjusted p=0.049) (Figure 2E), whereas Shannon diversity remained stable (linear mixed-model, β=0.02, FDR-adjusted p=0.62). Additional factors, including infant feeding mode (linear mixed-model, β=46.76, FDR-adjusted p=0.026) and the presence of pets at home (linear mixed-model, β=51.81, FDR-adjusted p=0.016), were also positively associated with plasmid richness (Figure S3F, S3G) (Table S6), with the effect of pets no longer significant after adjusting for infant feeding. Conversely, we did not observe a significant effect of time on plasmid composition (linear mixed-model, FDR-adjusted p=0.51) (Figure S3H). When assessing differences related to antibiotic timing at birth, we found that the overall plasmid composition was significantly different between the groups exposed and non-exposed to antibiotics (linear mixed-model, β=0.37, FDR-adjusted p=0.049) (Figure S3I) (Table S6). These results remained significant after correction for infant birth weight, which also showed a significant effect (linear mixed-model, β=-6e-4, FDR-adjusted p=0.015). To further estimate the change in the trajectories of plasmid composition in each infant over time, we used TCAM, a dimensionality reduction method for longitudinal ’omics data analysis.^35^ Using TCAM, we confirmed the nominally significant effects of both infant birth weight (Permutational Multivariate Analysis of Variance (PERMANOVA), 10000 permutations, R^2^=0.052, p=0.01) (Figure S3J) and timing of antibiotic administration (PERMANOVA adjusted for infant birth weight, 10000 permutations, R^2^=0.044, p=0.049) (Figure 2F). Lastly, we associated individual PTU abundances with available metadata. This identified 16 significant associations with infant age, including positive associations with two mobilizable PTUs (IMG_circular_plasmid_1528 and Circular_plasmid_286). Additionally, we identified two persistent PTUs associated with living environment (Plasmid_fragment_446) and maternal pre-pregnancy BMI (Plasmid_fragment_1422) (Table S7). Altogether, our results indicate subtle temporal changes in the infant gut plasmidome, with feeding mode, antibiotic timing at birth, and infant birth weight significantly impacting early gut plasmid diversity and community composition.

### Early infant gut plasmids represent expanding vectors of antibiotic resistance over time

Gut plasmids and other MGEs serve as reservoirs of ARGs and facilitate their transmission across gut microbial species.^9,13–15^ To understand the biological significance and prevalence of MGE-mediated ARG spread in the early infant gut, we screened and quantified the presence of ARGs in gut phages and plasmids. Plasmid genome annotation revealed 132 ARGs across early infant gut PTUs (weeks 1–6), while only 11 ARGs were found to be present in infant vOTUs, consistent with earlier reports.^36^ In total, almost 5% of infant gut PTU representatives had at least one ARG in their genome (75 PTUs). ARGs represented a significantly higher proportion of plasmid gene content (0.32%) compared to phage genomes (0.02%) (Chi-square test, p=5.15e-29) (Figure 3A). Given the lack of statistical power to perform the association analysis of ARGs in phages, we focused our analysis on the plasmid community in the early infant gut.

**Figure 3.**
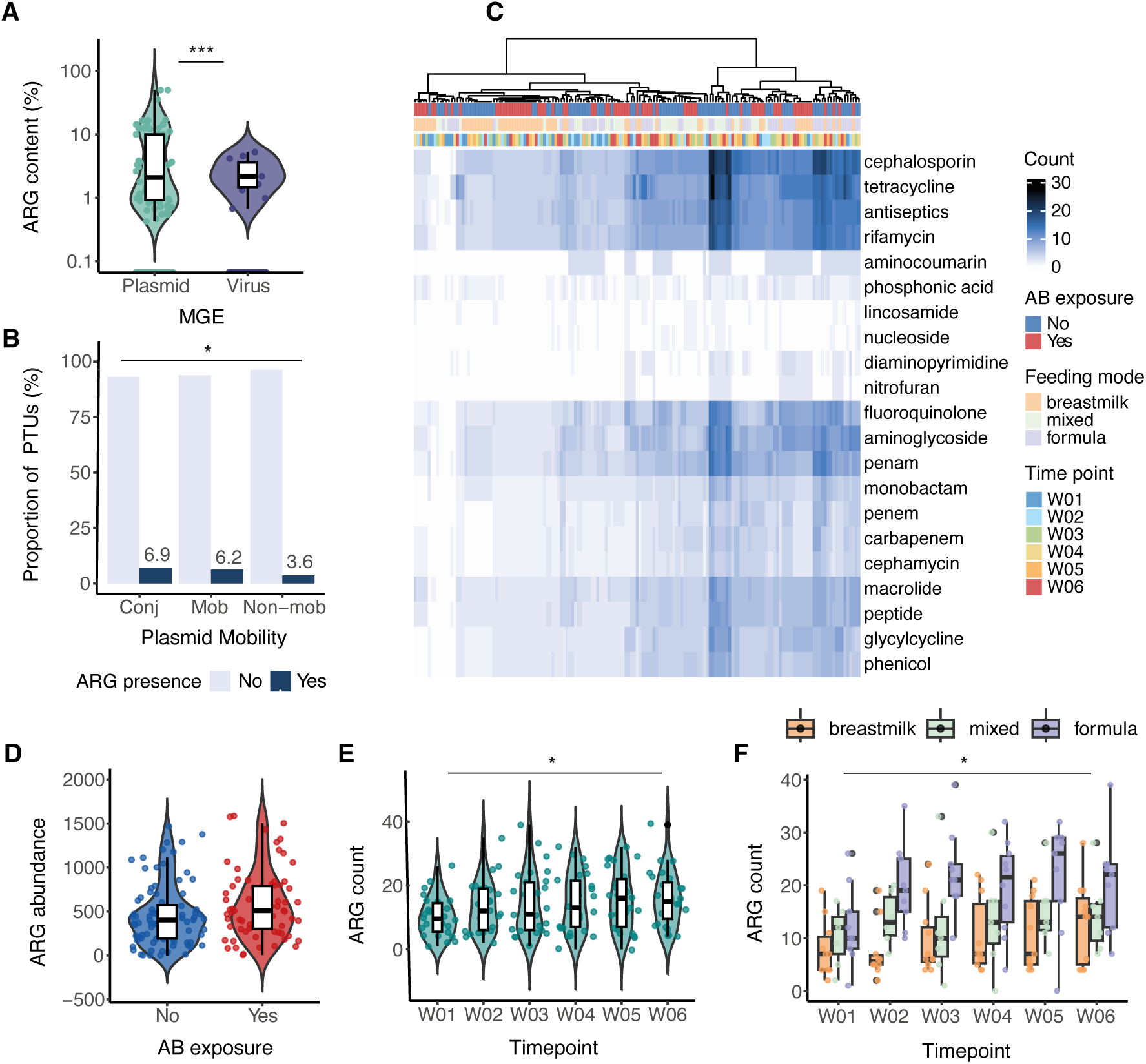
Early infant gut plasmids constitute expanding vectors of antibiotic resistance over time. (A) Boxplots and violin plots displaying the antibiotic resistance gene (ARG) content, estimated as the proportion of MGE-encoded genes identified as ARGs in PTU (green) and vOTU (purple) representative genomes. The y-axis is shown in log-scale. (B) Bar plot showing the proportion of PTUs with ARGs according to the plasmid mobility: conjugative (Conj), mobilizable (Mob), and non-mobilizable (Non-mob). Colours indicate the absence (light blue) or presence (dark blue) of ARGs. (C) Heatmap showing the number of ARGs identified per sample, aggregated by the antibiotic class to which they confer resistance. Dark blue indicates a higher number of ARGs. Lighter colours indicate a lower number. The top bars represent various factors: exposure to prophylactic antibiotics at birth (blue for non-exposed, red for exposed), feeding mode (orange for breastfeeding, light green for mixed-feeding, purple for formula-feeding), and timepoint. A dendrogram showing the hierarchical clustering of samples is displayed. (D) ARG abundance in PTUs based on exposure to prophylactic antibiotics at birth. (E) Number of ARGs in infant samples over the first 6 weeks of life. (F) Number of ARGs per time point grouped by infant feeding mode. Box plots illustrate the median values (middle line), interquartile range (box boundaries), and 1.5 times the interquartile range (whiskers).

The presence of ARGs in infant PTUs was also associated with the plasmid potential for mobilization, with mobilizable and conjugative plasmids showing increased presence of ARGs (Chi-square test, p=0.04) (Figure 3B). Further, we compared the load of plasmid ARGs between infants from both antibiotic treatment groups and explored associations with additional metadata. For this, total ARG counts and abundances were estimated for each sample and categorized based on the specific antibiotic they confer resistance to and the corresponding drug class. For instance, clustering by antimicrobial classes revealed cephalosporin and tetracycline to have the highest number of known resistance genes in the infant plasmids (Figure 3C). Overall, we found no statistically significant differences in the total number of ARGs (linear mixed-model, β=-3.83, p=0.11) or in their abundance (linear mixed-model, β=0.46, p=0.16) between infants exposed and not exposed to antibiotics at birth, although there was a trend towards increased abundance among exposed infants (Figure 3D). Similarly, we did not detect a significant effect of antibiotic exposure at birth on individual antibiotic resistance features after multiple-testing correction, although several nominally significant associations were observed (Tables S8, S9). However, we did find a significant increase in the number of ARGs detected in PTUs over time in infants (linear mixed-model, β=1.009, FDR-adjusted p=0.014) (Figure 3E) (Table S9). This increase was also observed in association with formula-feeding (linear mixed-model, β=9.38, FDR-adjusted p=0.018) (Figure 3F), driven by the increase in PTU richness. Taken together, our results suggest that early-life gut plasmids constitute an important reservoir of ARGs that increases over time and is modulated by infant feeding mode. In contrast, ARG prevalence remains low in early phage colonizers.

### Maternal–infant MGE strain-sharing is rare in the first 6 weeks of life and is influenced by infant feeding mode

To further characterize the maternal effect on the infant gut mobilome, we explored strain-level similarity and the potential transmission of viruses and plasmids from mothers to their infants in all available samples (including infant samples from months 6 and 12). For this, we selected vOTUs and PTUs present in at least two metagenomes and compared their strain-level profiles between each pair of samples with sufficient coverage and genome overlap (see Methods). Strain-level profiling of PTUs resulted in 40,324 pairwise comparisons involving 1,333 PTUs. This number was notably lower for viral strains (4,828 pairwise comparisons involving 613 vOTUs), indicative of the higher individual-specificity of vOTUs. Nonetheless, comparison of the maternal–infant distances of both plasmid (n=3,358) and phage (n=165) strains revealed a significantly higher strain similarity in related pairs (Permutation-based p<0.001) (Figure 4A, 4B).

**Figure 4.**
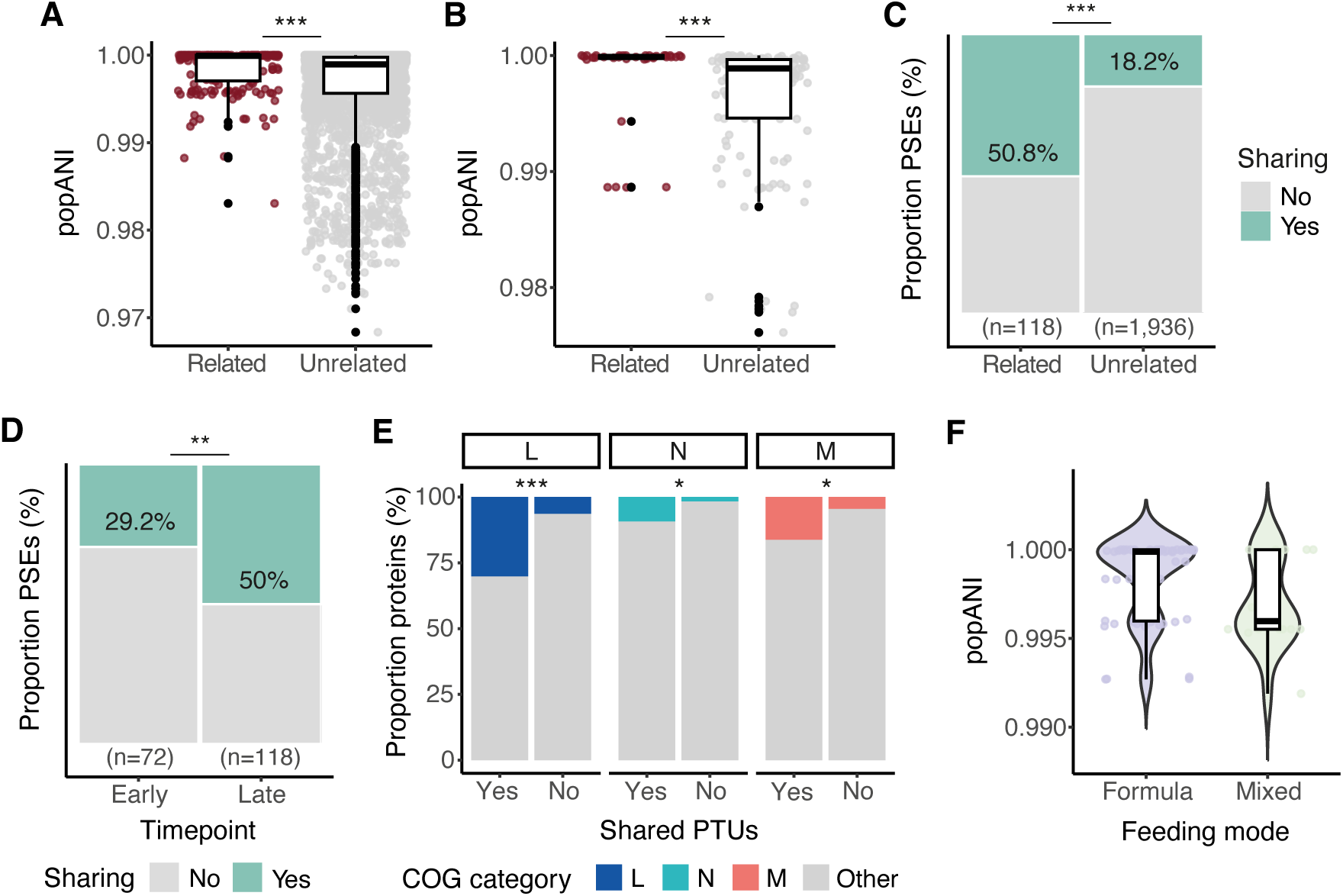
Maternal–infant MGE strain-sharing is rare in the first 6 weeks of life and influenced by infant feeding mode. (A,B) Box plots displaying the population average nucleotide identity (popANI) across (A) plasmid strains within each PTU and (B) viral strains within each vOTU, based on mother–infant relatedness. Median values (central line), interquartile range (box boundaries), and 1.5 times the interquartile range (whiskers) are shown. Comparisons between mother–infant pairs are shown in dark red. Non-pairs are shown in grey. (C,D) Mosaic plots showing the proportion of PTU strain comparisons identified as potential sharing events (PSEs) according to (C) mother–infant relatedness and (D) infant sample timepoints, categorized as early (weeks 1–6) or late (months 6 and 12). (E) Stacked barplots showing the COG functional categories enriched in shared PTUs. Categories include L: Replication, recombination, and repair (dark blue), N: Cell motility (light blue), and M: Cell wall/membrane/envelope biogenesis (red). All remaining functional categories are grouped into ’Other’ and displayed in grey. (F) Boxplots and violin plots illustrating the distribution of popANI values for plasmid strain comparisons within each PTU, categorized by infant feeding mode (purple for formula-feeding and light green for mixed-feeding).

To elucidate a potential maternal source of the early infant gut mobilome, we next explored potential viral and plasmid strain-sharing events in mother–infant pairs. For this, we applied the previously established cut-off of 99.999% population average nucleotide identity (popANI) for both phages and plasmids, defined based on the analysis of subspecies comparisons within the same infant over time.^37^ When considering all infant timepoints, we observed a significant difference in the proportion of plasmid-sharing events between related and unrelated mother–infant dyads (Chi-square test, p=2.12e-17) (Figure 4C). In total, we found 60 plasmid-sharing events across related pairs, involving 48 different PTUs. Notably, plasmid-sharing was significantly enriched at the later month 6 and month 12 timepoints (Chi-square test, p=7.58e-3) (Figure 4D), suggesting alternative MGE-sharing routes beyond vertical maternal–infant transmission. Plasmid circularity (Chi-square test, p=0.69) and potential for mobilization (Chi-square test, p=0.18) were not associated with different strain-sharing rates among related pairs (Figure S4A, S4B). Despite this, 45.8% (n=22) of the shared PTUs were mobilizable, and several exhibited potential transmission from week 1 of the infant’s life (Figure S4C). Functional annotation of plasmid genomes revealed that shared PTUs showed an enrichment of genes involved in replication, recombination, and repair; cell wall/membrane/envelope biogenesis; and cell motility (Fisher’s exact test, FDR-adjusted p=2.46e-5, 0.019, and 0.025, respectively) (Figure 4E). However, we found no ARGs to be encoded by any shared PTU. Analysis of viral genomes revealed similar results, with a significantly higher strain-sharing rate in related mother–infant pairs only when including comparisons from later months (Chi-square test, p=7.38e-9) (Figure S4D). In fact, all 12 potential sharing events in related pairs involving phage strains occurred after 6 months of infant age, with *Bacteroides*, *Alistipes*, and *Bifidobacterium* as the predicted viral hosts.

We next hypothesized that infant feeding mode could affect maternal–infant gut MGE-sharing in the first weeks of life. Our plasmid strain-profiling of mother–infant pairs (n comparisons=72, excluding months 6 and 12) found no common PTUs with sufficient genome coverage and overlap in the breastfed group, indicating highly distinct mother– infant gut plasmid profiles. On the contrary, most strain comparisons involved pairs using formula as infant feeding source (n=56). The statistical analysis comparing plasmid strain genome similarity (Wilcoxon rank sum test, p=0.08) and the proportion of sharing events (Fisher’s exact test, p=1) in the mixed-fed and formula-fed groups did not yield significant results, although we observed a trend towards increased similarity in pairs using formula as the infant feeding source (Figure 4F). All these findings indicate a potential route for maternal–infant transmission of gut MGEs. Strain-sharing is more prevalent for plasmids, is influenced by infant feeding mode during the early weeks, and most commonly occurs through horizontal transmission or environmental strain co-acquisition after the infant reaches 6 months of age.

## Discussion

MGEs, such as viruses and plasmids, play a crucial role in shaping gut microbial evolution and fitness through HGT.^4–12^ However, our understanding of their dynamic behaviour in early life and their contribution to the initial spread of ARGs remains limited. Here, we used an assembly-based approach to reconstruct the gut viral and plasmid communities during the first 6 weeks of life in infants born by CS. We explored their composition and temporal dynamics, which were modulated by intrinsic and extrinsic factors, and their role in disseminating ARGs within the gut microbial community. Furthermore, the inclusion of maternal samples collected around birth enabled us to investigate the potential maternal origin of early-life vOTUs and PTUs at strain-level.

Our study generated a comprehensive gut viral catalogue, significantly expanding the known phage diversity in early infancy. Although previous studies have explored longitudinal changes in the early-life gut virome,^17,19,21^ none have employed such dense sampling during the initial weeks of infant life. Our study design allowed us to reveal that the gut virome exhibits minimal diversity and compositional changes in the first 6 weeks of life, with only a slight increase in the number of detected viral species observed over time. This suggests that the significant changes in the infant gut virome primarily occur in later months, likely driven by factors such as increasing environmental exposures and dietary factors, which are known to have a profound impact on infant gut bacterial communities.^26,38,39^ In agreement with previous research, our analysis highlighted the significant individual-specificity of the gut phageome,^19,21,40^ with almost 90% of the species unique to each infant. This individual variation contrasted with the high temporal stability and persistence observed in the first weeks of life, aligning with previously reported findings in both infants and adults.^19,40^ Previous studies have suggested that DGRs may contribute to phage persistence in the gut by facilitating bacterial host switching.^20,34^ Contrary to this hypothesis, however, we did not find an enrichment of DGR systems in the persistent vOTU genomes, suggesting that the impact of these systems on viral persistence may only begin at later stages of infant life.

To provide a more comprehensive view of early-life gut MGEs, we aimed to characterize plasmid genomes in our samples. Characterization of plasmids presents significant challenges due to the complex nature of plasmids in metagenomes and the difficulties in differentiating them from other MGEs.^41,42^ To address these limitations, we employed a combined approach that used specific plasmid assemblers to reconstruct circular plasmids, alongside the identification of plasmid sequences from metagenomic contigs. Clustering plasmids into PTUs represents an additional challenge due to frequent HGT events, which rapidly reshape their genomes. While some methods rely only on backbone sequences to cluster plasmids,^42^ we employed a whole-sequence ANI-based approach to ensure highly similar gene content within each cluster, including ARGs, across entire plasmid genomes. By using this approach, we generated an extensive gut plasmid catalogue that comprised approximately 2,000 different PTUs in infant samples. Consistent with the patterns in viral richness, plasmid richness increased over the first 6 weeks of life, while the overall composition remained stable. Further classification of identified PTUs based on their capacity for mobilization revealed that a significant proportion of early-life PTUs encoded either mobilization genes or complete conjugation systems, indicating their ability to horizontally transfer the genetic material among early gut bacteria. Remarkably, we observed a significant association between plasmid mobilization potential and their abundance, prevalence, and persistence in the early infant gut, suggesting that plasmids with higher conjugative ability are more likely to establish and maintain their presence in the gut microbiota during early development.^23,43^

We identified several factors associated with vOTU and PTU diversity and community composition. Notably, infant feeding mode significantly impacted the richness of both vOTUs and PTUs, with formula-fed infants exhibiting higher richness than breastfed infants, even after controlling for infant age. At overall community composition level, no factors were observed to significantly alter the gut virome composition at this stage of life, while the early gut plasmid composition was significantly affected by both antibiotic timing at birth and infant birth weight. This aligns with earlier studies that have reported changes in early-life gut mobilome development related to gestational age and birth weight,^44,45^ as well as the reported impact of antibiotic treatment on plasmid abundances.^46^ Analysis of vOTU abundances aggregated by predicted viral hosts revealed several significant associations, but these were primarily driven by changes in the abundances of the bacterial hosts rather than by independent changes in the vOTUs. For instance, the positive associations of *Lacticaseibacillus* and *Clostridium* phages with formula-feeding did not persist after correcting for host abundance. These bacterial species have previously been observed to increase in abundance with formula-milk consumption, and they are used as probiotics in infant formula.^47–50^

The first weeks of infant life represent a critical window known to shape immune development in the newborn, potentially influencing the risk of chronic conditions later in life.^1–3^ Therefore, understanding how MGEs may contribute to the spread of ARGs is particularly relevant. We found that initial gut plasmids constitute an important reservoir of ARGs, with approximately 5% of total infant gut PTUs carrying predicted ARGs in their genomes. Importantly, the ARG load was also associated with plasmid mobilization potential, as mobilizable and conjugative PTUs displayed an increased presence of ARGs. In contrast, our findings suggest that early phage colonizers play a limited role in the spread of ARGs, which agrees with previous reports.^36^ While it has been hypothesized that antibiotic exposure at birth or in the early postnatal period impacts the early-life resistome,^51,52^ we did not observe a significant increase in ARG load in the MGEs of infants exposed to prophylactic antibiotics during CS. This indicates that the observed effect of antibiotic exposure on plasmid composition is not strongly associated with higher ARG content.

Thus far, the origin of the early infant gut mobilome remains largely unexplored.^31,32^ The limited number of birth cohorts that include maternal sample collections hampers our understanding of the initial establishment and origin of infant MGEs, as well as limiting exploration of possible maternal–infant colonization routes. Strain-resolved analyses are essential to determine the potential transmission of MGEs. Previously, we had demonstrated potential co-transmission of viral and bacterial strains from mothers to infants using infant samples collected from ages ranging 1 to 12 months.^19^ Supporting these findings, our current results indicate a higher MGE strain genome similarity and sharing rate in related mother–infant pairs. Specifically, we observe that plasmid strains are more commonly shared in mother–infant pairs than viral strains. Importantly, we observed that the plasmid strain-sharing rate was not associated with the plasmid’s potential for mobilization. In addition, most potential sharing events seemed to occur between 6 and 12 months of infant life, suggesting that the transmission of MGEs between mother and infant primarily derives from horizontal transmission after 6 months of age. This aligns with studies indicating limited maternal microbial transmission at birth in CS-born infants.^53–55^ Additionally, MGE strain-sharing may also result from the co-acquisition of strains from a common environmental source, rather than solely through direct mother-to-infant transmission.^53,56–58^ Among the mother–infant shared PTUs, various biological functions were enriched, but no encoded ARGs were detected. However, due to the limited sample size and the relatively low number of sharing events identified, further investigation is needed to accurately determine the potential for direct maternal–infant transmission of ARGs through MGEs. Early-life feeding mode has also been proposed as a potential source of MGEs for the developing infant gut.^29,32,59^ Our MGE strain comparison analysis did not identify common PTUs or vOTUs in mother– infant pairs using breast milk as the infant feeding source in the first 6 weeks of life, with all the potential sharing events detected in formula-fed infants. Thus, our findings suggest a potential protective role of breastfeeding against the early colonization of the immature infant gut by maternal gut MGEs. Expanding the research to larger cohorts, including vaginally born infants, will be essential to determine if these findings can be consistently replicated.

In summary, our study provides a detailed examination of MGEs in the early-life gut ecosystem of infants born by CS. By reconstructing and analysing gut viral and plasmid communities during the first 6 weeks of life, we observed minimal changes in the diversity and composition of MGEs over this period and determined their high individual-specificity and persistence in the gut. We found that early-life gut plasmids constitute an important reservoir of ARGs, with antibiotic exposure at birth and birth weight significantly influencing plasmid composition. Additionally, our findings suggest a potential maternal–infant transmission route of MGEs that may occur primarily through horizontal transfer in the later months of the first year of life. This comprehensive analysis sheds light on the dynamic behaviour of MGEs and underscores the need for further research to explore their interaction with gut microbes and their long-term implications.

## Acknowledgements

The authors would like to thank all patients for their participation in the trial. We thank Jelmer R. Prins, Sicco Scherjon and Marloes Kruk for assisting with the data collection and processing. We thank Kate Mc Intyre for editing our manuscript. We also thank the Genomics Coordination Center and the Center for Information Technology of the University of Groningen for providing access to the Gearshift and Hábrók high-performance computing clusters. A.F.P. was awarded a De Cocks-Hadders Stichting grant (2023-50). T.S. holds a scholarship from the Junior Scientific Masterclass, University of Groningen, and holds a De Cocks-Hadders Stichting grant (Winston Bakker Fonds WB-08). S.G. was awarded a De Cock-Hadders Stitching grant (2021-08). The work of S.R. is conducted at the U.S. Department of Energy Joint Genome Institute (https://ror.org/04xm1d337), a DOE Office of Science User Facility, supported by the Office of Science of the U.S. Department of Energy operated under Contract No. DE-AC02-05CH11231. J.F. is supported by the Netherlands Organization for Scientific Research (NWO) Gravitation grant Netherlands Organ-on-Chip Initiative 024.003.001, European Research Council (ERC) Consolidator grant 101001678 and NWO VICI grant VI.C.202.022. A.Z. is supported by the ERC Starting Grant 715772, NWO VIDI grant 016.178.056, EU Horizon Europe Program grant INITIALISE (101094099) and NWO Gravitation grant Exposome-NL (024.004.017).

## Author contributions

A.F.P., T.S. and A.Z. conceived the project. T.S. and A.Z. designed the randomized controlled trial. T.S. wrote the trial protocol. T.S. and A.Z. acquired funding for the trial and experiments. T.S. carried out the randomized controlled trial together with the research team of the Department of Obstetrics & Gynecology. T.S. performed the lab experiments. A.F.P. performed the bioinformatic analysis, under the supervision of S.G., A.G. and S.R. N.K. collaborated in the host prediction and functional annotation analysis. A.F.P. carried out the statistical analysis supervised by A.K. A.F.P., T.S., S.G., J.F., A.K. and A.Z. were involved in data interpretation. A.F.P. wrote the manuscript. All authors discussed the data and assisted in writing the manuscript. All authors have read and agreed to the published version of the manuscript.

## Methods

### Lead contact

Requests for further information and resources, software, reagents, and data sharing should be directed to the lead contact: Alexandra Zhernakova

### Materials availability

This study did not generate new unique reagents.

### Data and code availability

Sample metadata and quality-trimmed sequencing reads have been deposited at the European Genome-Phenome Archive (study ID: EGAS00001007571) and are publicly available as of the date of publication. Accession numbers are listed in the key resources table. All original code has been deposited on Github and is publicly available as of the date of publication.

https://github.com/GRONINGEN-MICROBIOME-CENTRE/CS_BABY_BIOME/tree/main/MOBILOME

### Study design and population

The CS Baby Biome RCT was conducted at the University Medical Center Groningen (UMCG), a tertiary referral centre, in the Departments of Obstetrics and Gynecology and Genetics. Recruitment occurred between 2019 and 2022. The study protocol (CCMO: NL61493.042.17) was approved by the UMCG’s Medical Ethics Committee (2017/240) and written informed consent was obtained from all parents. The study was registered as NCT06030713 on ClinicalTrials.gov: https://clinicaltrials.gov/ct2/show/NCT04066023.

Study design, participant recruitment, and randomization procedures are described in our previous publication.^30^

### Metadata collection

Information regarding CS indication, pre-pregnancy BMI, weight gain, gestational age, and infant characteristics were extracted from hospital records and stored in REDCap.^60^ Weekly digital questionnaires from parents covered feeding mode, medication use, environmental factors (exposure to pets and living environment), and infant health.

Feeding mode information was supplemented by patient records. Early-life feeding mode was calculated as the most frequent mode in the initial 6 weeks.

### Sample collection

Meconium samples were collected from neonates at the maternity ward and immediately transported to the UMCG (Department of Genetics) for storage at -20°C. Subsequent stool samples were collected from infants at home by their parents at weeks 1–6. Parents were provided with written and oral instructions on sample collection and storage by the research nurse. Collected stool samples were immediately frozen at -20°C in the home freezer.

In addition, maternal stool samples were collected from mothers shortly after birth and stored at -20°C at the Department of Genetics. At 6 weeks post-partum, maternal stool samples were collected, transported under frozen conditions, and stored at -80°C until DNA extraction. The same collection and storage procedures were followed for infant stool samples at 6 and 12 months post-partum. Umbilical cord blood was also collected to determine the infant’s exposure to cefazolin, as previously described.^30^

### Sample processing and *de novo* assembly

DNA extraction, library preparation, and shotgun metagenomics sequencing were performed as described in our earlier publication.^30^ The shotgun metagenomic sequencing reads were subjected to quality control, including the removal of human DNA contamination (reference human genome GRCh37), adaptor, and low-quality sequence trimming using BBDuk (v39.01) (http://bbtools.jgi.doe.gov) and Kneaddata (v0.10.0).^61^ Quality-filtered and trimmed reads from each sample (including unmatched reads) were assembled into contigs using MetaSPAdes (v3.15.5).^62^

### Bacterial abundance profiling and binning

Quality-trimmed reads were used to estimate the taxonomic composition of metagenomic samples using the MetaPhlAn4 tool with the MetaPhlAn database of marker genes mpa_vOct2022 and the ChocoPhlAn species-level genomic bin database (vOct2022).^63^ Bacterial binning was performed separately for each metagenome using metaWRAP (v1.3.2).^64^ Bins were then dereplicated at 98% whole-genome average nucleotide identity (gANI) with dRep (v3.4.2),^65^ selecting metagenome assembled genomes (MAGs) with a minimum completeness of 75% and maximum contamination of 10%.

### Viral identification workflow

Metagenome assembled contigs ≥10 kb in length were analysed by VirSorter2 (v2.2.4),^66^ DeepVirFinder (v1.0),^67^ geNomad (v1.5.1),^68^ VIBRANT (v1.2.1),^69^ and Cenote-Taker3 (v3.3.0) (https://github.com/mtisza1/Cenote-Taker3). Different inclusion criteria were applied for each tool: viral score >0.5 (VirSorter2), viral score ≥0.94 (DeepVirFinder), viral origin prediction with ’virion’ database option (Cenote-Taker3), viral origin prediction with --disable-find-proviruses option (geNomad), and viral origin prediction with default parameters (VIBRANT). All predicted viral contigs were combined and processed with COBRA (v1.2.3) to improve their genome assembly contiguity and completeness.^70^ To ensure the viral origin and trim potential flanking host regions from all extended sequences, we applied the geNomad default end-to-end pipeline. Next, CheckV (v1.0.1)^71^ was used to: A) further trim predicted viral sequences to remove potential host regions using the ‘contamination’ module and B) assess genome quality of resulting sequences using the end-to-end pipeline and select contigs that met the following criteria: ≥50% completeness, viral/host gene ratio >1, and genome length <1.5 times the expected genome length. Predicted viral sequences were then clustered into vOTUs with viral genomes from several public databases, including MGV (n=189,680 genomes with >50% estimated completeness),^33^ GPD (n=82,621 genomes with >50% estimated completeness),^72^ IMG/VR (n=36,064 genomes with >50% estimated completeness belonging to *Duplodnaviria* realm and found in human-associated samples),^73^ ELGV (n=21,295 genomes with >50% estimated completeness), Viral RefSeq (n=5,199 genomes >10 kb after excluding *Riboviria* realm),^21^ Shah et al. (n=4,627 dsDNA viruses with >50% estimated completeness),^18^ Benler et al. (n=1,480 viral genomes belonging to *Uroviricota* phylum)^74^ and CrAss-like phage databases (genomes with >50% estimated completeness from Gulyaeva et al. (n=637)^75^ and Yutin et al. (n=648)^76^ and all viral genomes from Guerin et al. (n=249)^77^). Dereplication was performed using the MIUVIG recommended settings (95% ANI; 85% AF (aligned fraction)),^78^ using the CheckV scripts *anicalc.py* and *aniclust.py* with the ‘--min_ani 95 --min_tcov 85 --min_qcov’ parameters. The longest sequence in each cluster was selected as the vOTU representative. Genus- and family-level vOTUs were generated by clustering vOTU-representative viral genomes based on a combination of pairwise average AAI and gene-sharing profiles, following the approach of Nayfach et al.^33^

### Prediction of bacteriophage lifestyle

To predict the viral lifestyle (temperate or virulent) of the bacteriophages, we used BACPHLIP (v0.9.6),^79^ which predicts lifestyle based on the presence of "temperate-specific" protein domains. Phage genomes lacking lysogeny-associated protein domains are by default classified as “virulent”. As phage genomes in our dataset have different levels of completeness, we assigned the viral lifestyle based on criteria previously described.^80^ Phages predicted to be temperate (score >0.5) were assigned a “temperate” lifestyle and phages predicted to be virulent (score >0.5) were assigned a “virulent” lifestyle when their genome-completeness estimated by CheckV was ≥90%. A prediction of a virulent lifestyle for vOTUs with a completeness below this threshold was omitted based on the potential presence of lysogenic markers in the missing viral genomic regions. Eukaryotic viral genomes (belonging to the *Bamfordvirae* and *Shotokuvirae* kingdoms) were excluded from the prediction.

### Taxonomy assignment and DGR identification

Taxonomy assignment was performed using geNomad taxonomy module (v1.5.1),^68^ which provides taxonomy based on the alignment of the viral genes to a set of markers that cover most of the lineages recognized by the International Committee on Taxonomy of Viruses (ICTV). All vOTU-representative genomes were scanned for the presence of DGR systems as previously described,^81^ in three main steps: (A) identification of reverse transcriptases (RTs) in viral genes predicted by prodigal-gv (v2.11.0)^82^ based on matches to Hidden Markov Models (HMM) profiles using hmmsearch (v3.4)^83^, (B) detection of repeats around the identified RTs using blastn (v2.15.0)^84^ with options -word_size 8 -dust no -gapopen 6 -gapextend 2, and (C) selection of putative DGRs based on repeat patterns and RT length. DGRs with an adenine mutation rate <75% were excluded from further analysis.

### Viral host prediction

Genus-level host prediction of identified phages was performed using iPHOP framework (v1.3.3).^85^ Prediction was performed using the default database (iPHoP_db_Aug23_rw) and the default database enriched with dereplicated MAGs generated from the metagenomes of the present study. For each vOTU, the bacterial genus with the highest confidence score between both prediction approaches was selected as host. Host assignments were further validated by estimating the Spearman correlation between the vOTU abundance levels (RPKM) and the relative abundance of the assigned host.

### Plasmid identification workflow

Quality-checked metagenomic reads were assembled into circular plasmids using metaPlasmidSPAdes (v3.15.5)^86^ and SCAPP (v0.1.4)^87^ with *max_k*=55. Predicted circular sequences >1 kb in length were screened using geNomad (v1.5.1)^68^ with the *-- enable-score-calibration* option to retain only contigs of plasmid origin with an FDR < 0.05. In parallel, metagenomic assemblies generated using MetaSPAdes (v3.15.5)^62^ were also analysed to identify potential plasmid genomes using geNomad. Predicted plasmid sequences were classified as follows. Circular plasmids were sequences meeting the criteria of length >1 kb, FDR < 0.05 (estimated using *--enable-score-calibration)*, and the presence of direct terminal repeats (DTRs). Plasmid fragments were sequences with length >1 kb, FDR < 0.05, no DTRs, and a minimum of one plasmid hallmark gene. All predicted circular plasmid sequences were combined with complete plasmids from the IMG/PR database^88^ (defined by the presence of DTRs or by alignment to complete reference plasmids) identified from human metagenomic samples. Circular plasmids and plasmid fragments were then dereplicated separately and clustered into PTUs using the CheckV scripts *anicalc.py* and *aniclust.py* with strict (*--min_ani 95 --min_tcov 90 -- min_qcov 90*) and loose (*--min_ani 90 --min_tcov 85 --min_qcov 0)* parameters, respectively. Both plasmid sets were then combined using the loose settings, filtering fragment PTUs that clustered together with circular PTUs.

### Determination of plasmid potential for mobilization

The presence of proteins that are components of conjugation systems (VirB4 T4SS ATPases, relaxases, T4CPs, and other T4SS components) were detected by using the CONJScan (v2.0.1)^89^ module of MacSyfinder (v2.1.1).^90^ Briefly, this method relies on the comparison of plasmid protein sequences against the CONJScan HMMs, verifying both the system composition and genetic organization. Only matches with an e-value ≤ 0.001 and a minimum 50% profile coverage were considered. Origins of transfer (oriTs) were detected as described by Camargo et al.^88^, using blastn against a database of previously identified oriT sequences, with a minimum target coverage of 50% required. In cases of overlapping oriTs, only the hit with the lowest e-value was retrieved. Plasmids were classified as conjugative (encoding complete conjugation systems based on the presence of all mandatory components defined in CONJScan models), mobilizable (lacking complete conjugation systems but encoding a relaxase or oriT), and non-mobilizable (plasmids that did not meet the criteria for the previous two categories).

### ARG and functional annotation

ARGs in both PTU- and vOTU-representative genomes were annotated using the Resistance Gene Identifier (RGI) (v6.0.3) against the CARD 2023 database.^91^ Only perfect or strict hits were considered. The abundance of an ARG in each sample was calculated as: sum (counts ARG per MGE * abundance MGE). CARD annotation results were then processed using a custom R script to estimate the number and abundance of ARGs categorized based on the antimicrobials to which they confer resistance and the corresponding drug class. PTU-representative genomes were further annotated using BAKTA (v1.9.2),^92^ and COG functional categories were retrieved for the functional enrichment analysis of transmitted PTUs.

### vOTU and PTU abundance estimation

To estimate the viral and plasmid abundances in each sample, we mapped the metagenomic reads, filtered and quality-trimmed as described above, to the database of vOTU- and PTU-representative sequences generated using Bowtie2 (v2.4.5).^93^ Breadth of genome coverage by sequencing reads was calculated using the BEDTools (v2.30.0) command coverage.^94^ Each vOTU and PTU were considered present in a sample if the mapped reads covered >75% of the genome length.

### Viral and plasmid strain transmission analysis

Sequence alignment mapping files were processed using inStrain (v1.9.0)^37^ “profile” with a minimum mapQ score of 0 and an insert size of 160. inStrain “compare” was then used to estimate the between-sample genome similarity among all reconstructed consensus sequences of vOTU- and PTU-representative genomes that were present in a minimum of two samples. Only regions with a minimum of 5x coverage were included in comparisons. Sample pairs with <75% comparable regions for phage and plasmid genomes were excluded from the analysis. Population-level ANI (popANI) values were used to compare similarity between strains belonging to the same vOTU or PTU. Phage or plasmid strains were considered as shared between samples if their genomic regions exhibited ≥99.999% popANI. Maternal–infant transmission was determined by comparing vOTUs and PTUs detected in ≥1 mother–infant pair.

### Statistical analysis

The association analysis with phenotypes was performed on early-life infant samples (week 1 to week 6) using linear mixed-models. Association analysis was performed on log-transformed RPKM abundances of vOTUs aggregated by bacterial host (≥5% prevalence), PTUs (≥10% prevalence), and ARGs (including ARG counts) grouped by the antimicrobials they confer resistance to (≥10% prevalence). vOTU and PTU diversity and composition variables were also used for association analysis with phenotypes. Associated phenotypes were infant age, birth weight, gestational age, maternal pre-pregnancy BMI, infant sex, infant feeding mode, living environment (city, village, farm), presence of cats/dogs at home and antibiotic exposure at birth. All models included the number of quality-trimmed reads and DNA concentration as fixed effects and individual infant ID as a random effect. Additional covariates, such as infant age, birthweight, feeding mode, or CLR-transformed bacterial host abundance, were included when specified. Statistical significance was determined at FDR < 0.05 using the Benjamini-Hochberg method. Alpha and beta diversity analysis of viral and plasmid communities were performed at vOTU- and PTU-level using R package *vegan* v2.6-4.^95^ Specifically, vOTU and PTU richness was calculated using the *specnumber()* function. Alpha diversity was estimated as the Shannon diversity index using the *diversity()* function. Beta diversity analysis was performed using Bray-Curtis dissimilarity calculated using the function *vegdist()*. Nonmetric multidimensional scaling (NMDS) was performed using the *metaMDS*() function from the *vegan* package. Two dimensions were generated to visualize the dissimilarity between samples, while a one-dimensional NMDS was utilized for analytical purposes. TCAM,^35^ a dimensionality reduction method for longitudinal ’omics data analysis, was used to combine multiple timepoints of the same individual into a single measure that represents the temporal trajectory of the individual. The resulting matrix was used for PERMANOVA via the *adonis2()* function in the *vegan* package. Using 1,000 permutations, we estimated the effect size (R²) and significance of phenotypes on vOTU and PTU composition based on Bray-Curtis dissimilarity. Statistical significance for various comparisons was determined using Fisher’s exact test (*fisher.test()*), Chi-square test (*chisq.test()*), and Wilcoxon rank sum test (*wilcox.test()*), all from the *stats* R package, as specified in the text. Significance of distance comparisons based on virome composition and strain genome similarity was assessed using a permutation test with 1,000 iterations. Post-hoc pairwise comparisons were conducted using Dunn’s test with the *dunn.test()* function from the dunn.test package.^96^ Two-sided statistical tests were used unless otherwise specified. All statistical analyses were performed and all plots were created using R (v4.2.2).^97^

