## Supplementary Figures for "Dynamics and Determinants of the Gut Mobilome in Early Life"

A. Fernández-Pato et al.

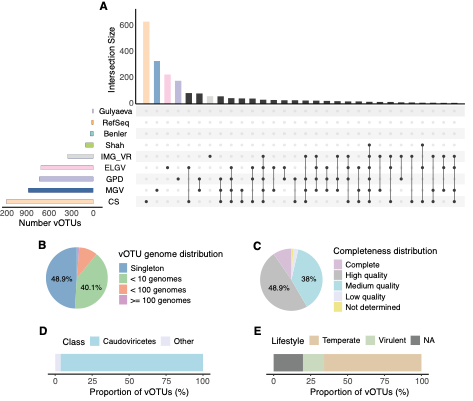

**Figure S1. Generation and characterization of the infant gut viral catalogue.**

(A) UpSet plot comparing the CS Baby Biome vOTU catalogue (n=2,263) against the public human gut virus databases MGV (n=189,680), GPD (n=82,621), IMG/VR (n=36,064), ELGV (n=21,295), RefSeq (n=5,199), Shah et al. (n=4,627), Benler et al. (n=1,480) and Gulyaeva et al. (n=637). The comparison was made only for quality-filtered virus sequences (see Methods). Vertical lines connected by dots indicate the vOTU overlap between databases. (B,C) Pie charts illustrating (B) the distribution of the number of viral genomes within identified vOTUs and (C) the viral completeness distribution of vOTU representatives as determined by CheckV. (D,E) Bar plots displaying the distributions of (D) the class-level taxonomic assignment and (E) the viral lifestyle prediction of identified vOTU representatives.

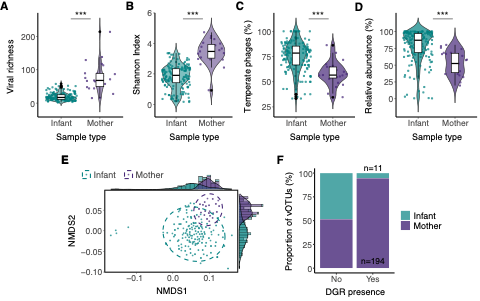

**Figure S2. Mothers and infants show a different gut virome diversity and composition.**

(A–D) The (A) viral richness, (B) alpha diversity (measured as Shannon Index), (C) proportion of temperate phages, and (D) relative abundance of temperate phages in infant (green) and maternal (purple) samples. In all cases, box plots show the median values (middle line), interquartile range (box boundaries), and 1.5 times the interquartile range (whiskers). (E) NMDS plot based on Bray-Curtis dissimilarity estimated on vOTU abundances. Each dot represents the viral composition of one sample, coloured according to maternal or infant origin. Ellipses represent 95% confidence regions, assuming a multivariate t-distribution of the data points. A density plot overlaid on a histogram for both NMDS dimensions is also shown. (F) Bar plot illustrating the proportion of vOTUs in which diversity-generating retroelements (DGRs) were detected coloured according to maternal or infant origin.

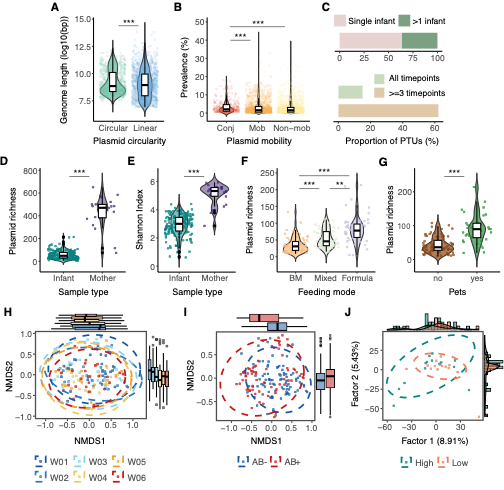

**Figure S3. Characterization of the early infant gut plasmid community, differences with maternal gut, and factors affecting its diversity and composition.**

(A) The genome length distribution (log10 scale) of circular versus linear PTUs. (B) Prevalence of PTUs categorized by mobility group: conjugative (Conj), mobilizable (Mob), and non-mobilizable (Non-mob). (C) Bar plots illustrating the proportion of PTUs found in a single infant (pink) versus multiple infants (green), and the proportion of PTUs present in at least three timepoints (light brown) or all timepoints (light green) within the same infant over the first 6 weeks of life. (D,E) The (D) PTU richness and (E) alpha diversity (measured as Shannon Index) comparison between infant and maternal samples. (F,G) PTU richness comparison according to (F) infant feeding mode (breast milk (BM), mixed-feeding, and formula-feeding) and (G) the presence of pets at home. (H,I) NMDS analysis based on Bray-Curtis dissimilarity estimated on PTU abundances. Each dot represents the viral composition of one sample, coloured according to (H) the timepoint and (I) antibiotic exposure at birth (blue for non-exposed, red for exposed). (J) Scatter plot of the leading TCAM factors estimated based on PTU abundances of infant samples. Each dot represents the PTU composition of a single infant over the first 6 weeks of life. Colours represent infant birth weight categorized into 'High' (dark green) and 'Low' (orange) based on the median value of 3.667 kg. In all cases, box plots show the median values (middle line), interquartile range (box boundaries), and 1.5 times the interquartile range (whiskers). Ellipses represent 95% confidence regions, assuming a multivariate t-distribution of the data points.

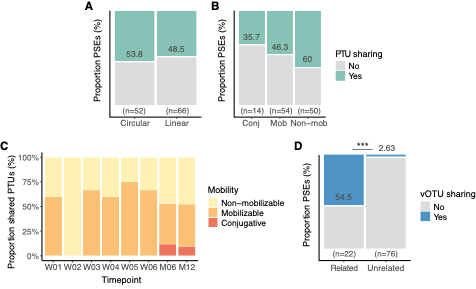

**Figure S4. Analysis of MGE strain-sharing according to plasmid circularity, mobilization potential, and mother**–**infant relatedness.**

(A,B) Mosaic plots displaying the proportion of PTU strain comparisons identified as potential sharing events (PSEs) based on (A) plasmid circularity and (B) mobility. (C) Bar plot showing the proportion of shared PTUs by plasmid mobility group (categorized as non-mobilizable (yellow), mobilizable (orange), and conjugative (red)) per infant sample timepoint. (D) Mosaic plot showing the proportion of viral strain PSEs based on mother–infant relatedness in all available samples.
